# Partial agonists and heteromeric Kainate receptor desensitization

**DOI:** 10.64898/2026.11.02.762920

**Authors:** Guadalupe Segura-Covarrubias, Ashlee Hoffman, Changping Zhou, Nami Tajima

**Author notes:** Correspondence to (N.T.) and (G.S-C.). Equal contribution (Guadalupe Segura-Covarrubias, Ashlee Hoffman, Changping Zhou).

## Abstract

Kainate receptors (KARs) are glutamate-gated ion channels that mediate fast excitatory synaptic transmission in the central nervous system. KARs assemble as homo- or heterotetramers composed of five subunits (GluK1–5), with heteromeric assemblies representing the predominant native population. Although KARs share overall structures and basic functional properties with other ionotropic glutamate receptor subfamilies, they exhibit distinct gating kinetics, particularly with respect to desensitization. While earlier studies have demonstrated how agonist binding drives gate opening and subsequent desensitization, the mechanisms governing partial agonism and desensitization of KARs, especially in heteromeric KARs, remain incompletely understood. Here, we combine electrophysiology and microscale thermophoresis binding assays to investigate how distinct KAR partial agonists control KAR desensitization and its recovery. While the onset rates of desensitization of homo- and heteromeric receptors activated by high/saturating concentrations of partial agonists are conserved between GluK2 and GluK2/GluK5 receptors, recovery from desensitization in heteromers is significantly slower than in homomers. Together with structural modeling and analysis of the GluK2 and GluK5 ligand binding domains (LBDs) in complex with orthosteric ligands, we show that agonist residence time and the extent of LBD bi-lobe closure across various agonists take precedence over subunit composition in governing KAR desensitization.

## Introduction

Kainate receptors (KARs) are a subfamily of ionotropic glutamate receptor (iGluRs) alongside alpha-amino-3-hydroxy-5-methyl-4-isoxazole propionic acid (AMPA), N-Methyl D-Aspartate (NMDA) and delta receptors^1^. KARs are ligand-gated ion channels that regulate synaptic plasticity and neuronal circuit through pre-, post-, and extra-synaptic mechanisms^2–5^, and are implicated in various neurological and psychiatric disorders, including epilepsy and schizophrenia^2,6^. They form homo- or heterotetramers comprising five subunits (GluK1–GluK5), with each subtype exhibiting distinct channel properties that expand KAR functional diversity. The low-agonist affinity subunit GluK1–3 can form functional homomeric and heteromeric receptors, whereas the high-affinity subunits (GluK4 and GluK5) act as obligate heteromeric subunits requiring co-assembly with GluK1–3 subunits. Each subunit consists of four domains; amino-terminal domain (ATD), the ligand-binding domain (LBD), the transmembrane domain (TMD) and C-terminal domain (CTD). The subtype diversity, together with alternative splicing^3,7,8^, RNA editing^9,10^, post-translational modifications^11–17^, and association with auxiliary and interacting proteins^18–27^, generates receptors with diverse pharmacological and channel properties. In the brain, native KARs predominantly assemble as heteromers^28,29^ with the GluK2/GluK5 subtype being the most abundant^30–32^. Structural and single-molecule imaging studies have established that heterotetrameric GluK2/GluK5 receptors assemble with a 2:2 stoichiometry in an alternating K2–K5–K2–K5 spatial arrangement, wherein GluK5 subunits occupy the pore-proximal (A/C) positions and GluK2 subunits are situated in the pore-distal (B/D) positions^33–35^. Specifically activating the high-agonist-affinity subunits in heteromeric receptors stabilizes them in intermediate conformations with only partial desensitization^35^. Functional studies have also established that homomeric and heteromeric KARs (e.g., GluK2 versus GluK2/GluK5) exhibit distinct gating kinetics, including desensitization, in addition to differences in ligand sensitivity^36–39^.

The KAR LBD possesses a bi-lobed, clamshell-like structure composed of an upper D1 lobe and a lower D2 lobe^40,41^. A variety of orthosteric ligands, including full and partial agonists as well as competitive antagonists, occupy overlapping binding sites within the LBD clamshell^42^. The agonist binding within the cleft between D1 and D2 induces clamshell closure, which in turn opens the ion channel formed by the TMDs, a conserved mechanism across all iGluRs, while competitive antagonists typically stabilize an open LBD conformation, maintaining receptors in inactive states^1,43,44^. Partial agonists are ligands that bind to and activate receptors but exhibit submaximal efficacy relative to full agonists^36^. At homomeric KARs, diverse partial agonists produce distinct peak responses, functional efficacies, and gating kinetics^45^. Extensive structural studies have characterized isolated KAR LBD conformations in complex with a variety of agonists and antagonists^42^. Additionally, we have previously shown that partial agonists stabilize full length GluK2 KAR in multiple intermediate/inactive conformations^46^, similar to observations in partial agonist-bound AMPA receptors^47,48^. Unlike other iGluRs, where the degree of LBD closure correlates directly with efficacy and agonist responsiveness^49–51^, the degree of KAR LBD clamshell closure alone does not fully predict agonist efficacy. Consequently, the mechanisms governing partial agonism at heteromeric KARs appear more complex^52^.

In this study, we investigate how partial agonist size and chemical structure dictate desensitization and recovery kinetics in heteromeric versus homomeric KARs. We first generated models of the LBDs of low- and high-affinity KAR subunits bound to partial agonists, compared their bi-lobe opening degrees with published ligand bound LBD structures, and determined the dissociation constant for each agonist. Next, we used patch-clamp electrophysiology to measure the gating kinetics of homomeric and heteromeric receptors activated by these agonists. This integrated approach allowed us to examine whether agonist-specific binding properties correlate with distinct gating kinetics in heteromers and to determine how these relationships are modified by the incorporation of high-affinity subunits.

## Results

### Structural Spectrum of GluK2 LBD Conformations and Interactions Across Full/Partial Agonists and Competitive Antagonists

To better understand how distinct ligands stabilize specific subunit conformations within low-affinity KARs, we first analyzed the conformation of the GluK2 LBD in complex with five ligands of varying steric bulk. This set comprised a full agonist (L-glutamate), three partial agonists of progressively higher molecular weight (MW) relative to glutamate [L-α-aminoadipic acid (L-AAA), kainate, and domoate], and a competitive antagonist comparable in size to domoate (UBP310) (Figure 1A). Because glutamate and L-AAA are highly flexible, containing three and four rotatable bonds in their core chains, respectively, with L-AAA exhibiting distinct conformations (Fig S1A), we performed physicochemical descriptor analysis and calculated their solvent-accessible surface area (SASA) and net change at pH 7.4. This analysis confirmed that all ligands possess similar predicted formal net charges at pH 7.4 (glutamate, L-AAA, kainate, UBP310: -1 e, whereas domoate: -2 e) (Fig. 1B), and their surface area at physiological pH strongly correlates with MW (Fig. S1B).

**Figure 1.**
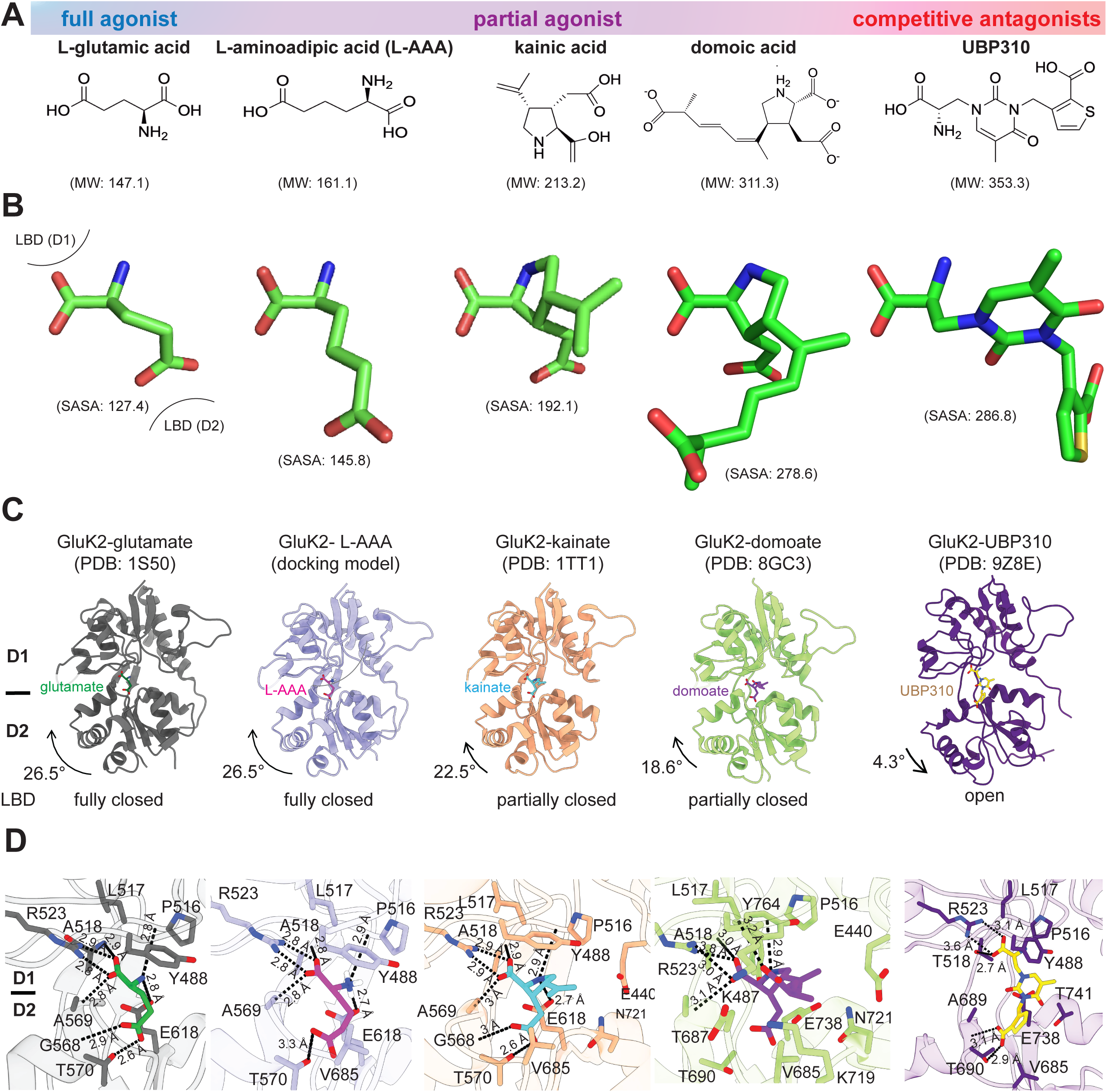
Chemical structures and ligand bound GluK2 LBD conformations. **(A)** 2D chemical structures, pharmacological classifications (full agonist, partial agonist, competitive antagonist), and molecular weights (MW, in g/mol) for L-glutamic acid, L-aminoadipic acid (L-AAA), kainic acid, domoic acid, and UBP310. **(B)** 3D stick representations of the ligands in their bound conformations showing LBD domain contact orientations (D1 top, D2 bottom) and calculated SASA values (Å^2^). **(C)** Structural models showing the degree of LBD domain closure induced by each ligand, including the rotation angle relative to the apo state (PDB 9C5Z). **(D)** Close-up views of the interactions between ligand and key amino acid residues within the GluK2 LBD ligand binding pocket.

Initially, we attempted to determine the structure of GluK2 in complex with L-AAA. However, the low affinity of the ligand required high ligand concentration (>10 mM) to maintain occupancy and prevent complex dissociation, introducing technical challenges. These included sample precipitation of purified full-length tetrameric GluK2 on cryo-electron microscopy (cryo-EM) grids and non-protein crystallization (salt or ligand) during X-ray crystallography trials with the isolated GluK2 LBD. To circumvent these experimental limitations, we generated an AlphaFold 3 model of the rat GluK2 LBD complexed with L-AAA using its Chemical Component Dictionary (CCD) code UN1 and compared it with published structures of GluK2 LBD bound to the other four ligands. Superimposing the L-AAA bound GluK2 LBD onto the glutamate bound structure (PDB 1S50)^40^ demonstrates that glutamate and L-AAA adopt similar binding modes (Fig. S1C, D). Based on its chemical structure and previously observed conformations in complexed with other L-AAA binding proteins^53,54^ (Fig. S1A), we reason that the low affinity of L-AAA is driven by the ligand conformational flexibility or instability of its binding pose within the agonist-binding pocket (Fig. S1E).

Comparison of these five orthosteric ligand-bound GluK2 LBD structures relative to the apo GluK2 (PDB 9C5Z)^46^ conformation illustrates the spectrum of rigid-body LBD bi-lobe closure (for agonists) and opening (for the competitive antagonist) induced by ligand binding (Fig. 1C). The glutamate bound GluK2 LBD structures solved by X-ray crystallography or cryo-EM (PDB 1S50 and 9MZJ)^40,55^ and the predicted L-AAA bound LBD are stabilized in the fully LBD bi-lobe closed conformation with 26.5° more closed than the apo state, whereas the reported kainate- and domoate-bound GluK2 LBD structures (PDB 1TT1 and 8GC3)^40,46^ showed partial closed conformations with closure angles of 22.5° and 18.6°^40,56^, respectively. In contrast, the competitive antagonist-bound GluK2 LBD adopts a 4.3° more open conformation (PDB 9Z8E)^57^ than the apo (PDB 9C5Z)^46,58^, even though UBP310 is comparable in size to domoate. We observe the spectrum of LBD spanning fully closed, partially closed, unliganded apo, and open state. As previously reported, smaller ligands (glutamate, L-AAA, and kainate) share conserved interactions across both the D1 and D2 lobes, while bulkier ligands such as domoate and UBP310 engage distinct residues especially on the D2 lobe to stabilize less closed LBD conformations (Fig. 1D).

### Distinct binding affinity of full and partial agonists

All tested ligands possess an α-amino acid motif that serves as the core binding group interacting with the D1 domain of the LBD, whereas distinct extending moieties engaging the D2 domain dictate binding affinity (Fig. 1B). To directly compare the binding affinities of these agonists on homomeric KARs, we expressed and purified the isolated GluK2 LBD construct to determine dissociation constants (Kd) via microscale thermophoresis (MST). Briefly, the GluK2 S1 and S2 segments were connected via a short GT linker and fused to an N-terminal hexahistidine-SUMO tag in a pET-22b expression vector. The fusion protein was overexpressed in E. coli, purified by Ni-NTA affinity chromatography, and cleaved with ULP-1 SUMO protease using a protocol modified from published NMDAR GluN1/GluN2B LBD purifications prior to His-tag labeling for MST analysis.

All three agonists showed concentration-dependent binding curves with substantially different apparent affinities (Fig 2A). Kainate displayed the highest affinity, with a Kd of 0.057 ± 0.0082 μM followed by glutamate (Kd: 0.42 ± 0.06 μM), while L-AAA exhibited a markedly higher Kd of 308.7 ± 46.0 μM (Fig. 2B). This low affinity is consistent with previous reports showing that high concentrations of L-AAA (10–40 mM) are required to activate GluK2^45^. This high Kd, 740-fold higher than that of glutamate, supports the concept that ligand structure and agonist-binding pocket dynamics strictly dictate ligand accommodation, which is critical for the gating and full activation of GluK2 KARs. Consequently, these MST measurements demonstrated pronounced ligand-dependent differences in binding affinity.

**Figure 2.**
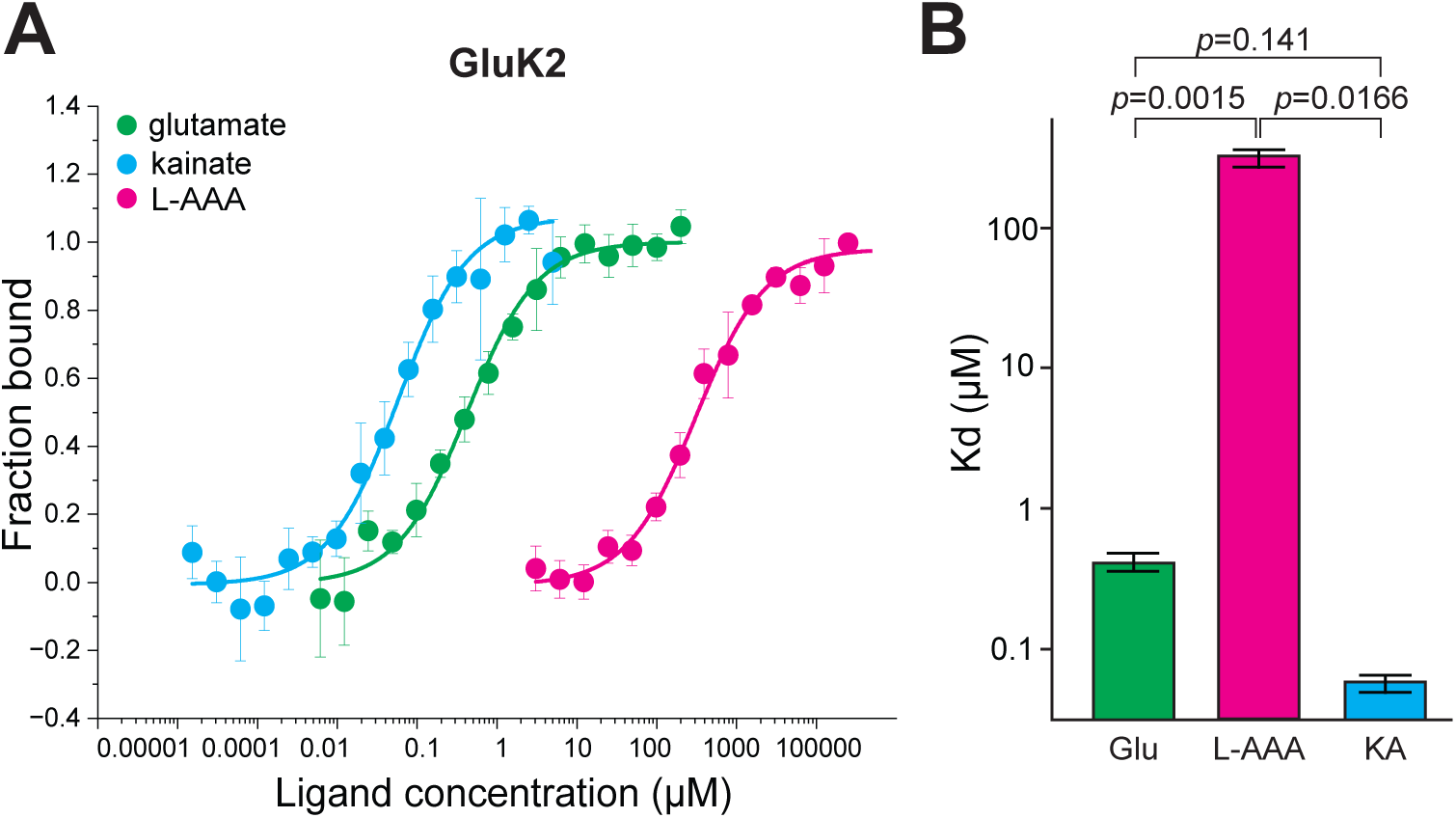
Binding affinities of agonists for GluK2. **(A)** Microscale thermophoresis (MST) binding curves for GluK2 in response to increasing concentrations of glutamate (Glu, green), L-α-aminoadipic acid (L-AAA, magenta), and kainate (KA, blue). Data points represent the fraction bound plotted against ligand concentration. Curves were fitted to a standard single-site binding model. **(B)** Summary bar graph comparing equilibrium dissociation constants (Kd) for glutamate, L-AAA, and kainate derived from MST measurements. Data are presented as mean ± SEM. Statistical significance was assessed using student’s t-test.

### Full- and partial agonists show distinct gating kinetics of GluK2 KAR

Having established that these four agonists display markedly different apparent affinities for the isolated GluK2 LBD, we next evaluated how ligand identity impacts homomeric GluK2 receptor kinetics, focusing specifically on desensitization and recovery from desensitization. We expressed wild-type rat GluK2 in HEK293T cells, activated the receptors by three distinct agonists, glutamate (10 mM), L-AAA (30 mM), and kainate (0.5 mM), and compared these responses to our previously published data of GluK2 elicited by domoate (10 μM)^46^. In GluK2 receptors, glutamate showed the fastest time constant of desensitisation (τ_desensitization_) (7.5 ± 0.3 ms, n=14), followed by kainate (10.6 ± 0.6 ms, n=8) and L-AAA (14.7 ± 2.1 ms, n=6) (Fig. 3A–B). Desensitization decay kinetics were uniformly fitted using a single exponential function. Despite large differences in Kd, desensitization decay rates varied by at most two-fold across all three agonists. τ_desensitization_ was significantly faster with glutamate than with kainate (p=0.0024) or L-AAA (p=0.0396), while the comparison between kainate and L-AAA showed no statistically significant difference (p=0.23) (Fig. 3A, B), a kinetic pattern that is similar but slightly differs from the previously reported behaviour^59^. We next evaluated how these agonists affect recovery from desensitization. While the high-affinity agonist kainate exhibited significantly slower recovery kinetics (3.2 ± 1.2 s, n=5) than glutamate (1.1 ± 0.3 s, n=11), L-AAA, which possesses a high dissociation constant (i.e., low-affinity), demonstrated markedly faster recovery (0.2 ± 0.1 s, n=6) than glutamate (Fig. 3C– D).

**Figure 3.**
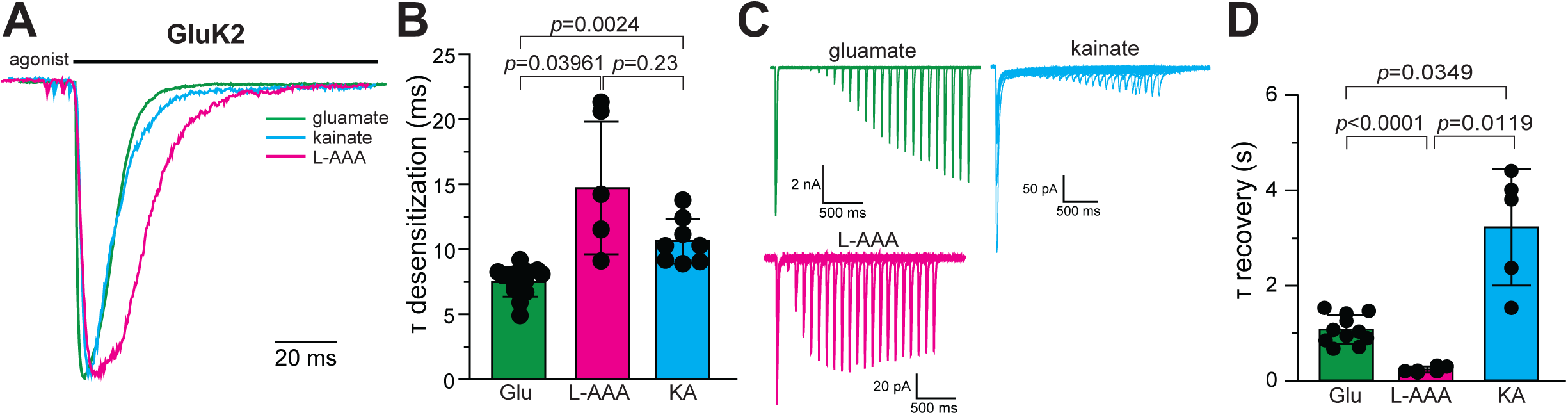
Desensitization and recovery kinetics of homomeric GluK2 receptors evoked by different agonists. **(A)** Representative whole-cell recordings from HEK293T cells expressing GluK2 activated by 10 mM glutamate, 30 mM L-AAA, or 0.5 mM kainate for 1 s (black bar). **(B)** Mean desensitization rate constants for GluK2. **(C)** Representative current traces of GluK2 activated by glutamate (green), L-AAA (magenta), and kainate (blue) during a two-pulse protocol, showing the interval between glutamate exposures ranging from 50 ms to 20 s. **(D)** Time constant (*τ*) of recovery from desensitization.

### Agonist predicted to induce similar conformational changes in GluK2 and GluK5 LBDs

Next, to assess subtype-dependent differences in KAR properties, we evaluated the LBD conformations and Kd values for these three agonists at high-affinity KAR subunits and compared the gating kinetics of heteromeric GluK2/GluK5 KARs with those of homomeric GluK2 receptors. Due to the lack of experimental structural data for L-AAA- and kainate-bound GluK5 LBD, we generated AlphaFold models and compared them to the apo-, glutamate-, and UBP310-bound GluK5 LBD structures we recently published^57^. Sequence identity between rat GluK4 and rat GluK5 LBDs is 84.4%, with kainate-binding residues within the agonist-binding pocket fully conserved (Figure S2B). In the modelled kainate-bound GluK5 structure, kainate adopts a conformation within the binding pocket superimposable onto the kainate-bound GluK4 LBD crystal structure^60^, validating the reliability of our computational model (Figure S2C). Comparison with the structure of GluK5 LBD in complex with the partial agonist α-amino-3-hydroxy-5-methyl-4-isoxazolepropionic acid (AMPA) (PDB 9OSI)^35^, another glutamate analogue that shares overlapping pharmacophores with kainate, shows a similar conformation of GluK5 LBD, with a root-mean-square deviation (RMSD) of 1.6 Å and the identical LBD bi-lobe opening degree to the predicted kainate-bound GluK5 LBD. The binding mode of L-AAA in GluK5 is predicted to resemble that in GluK2, mimicking glutamate binding in GluK5 with an identical degree of LBD domain closure as the glutamate-bound GluK5 (PDB 9Z8B)^57^ (Fig. 4A). Although the apo GluK2 and GluK5 display distinct LBD opening angles, structural comparison of these three agonist-bound conformations indicates that both full and partial agonists induce LBD closure to a similar extent as in GluK2. The glutamate (PDB 9Z8B)^57^ and L-AAA bound GluK5 LBDs showed 4°more closed conformation than the kainate-bound GluK5 LBD model (Fig. 4A). Because the agonist-binding pocket residues are identical (100% sequence identity) across all GluK1–5 subunits, these binding modes appear conserved between GluK2 and GluK5 (Fig. 4B). As observed in the glutamate and L-AAA bound GluK2 LBD structures, key ionic interactions, salt bridges, and hydrogen-bonding networks are conserved in GluK5. These include contacts with Arg506, Thr501, Ala499, Phe500, and Tyr572 on the D1 upper lobe, as well as Ser558, Thr553, and Glu607 on the D2 lower lobe. In contrast, kainate displays a distinct interaction pattern and fails to promote complete closure of the GluK5 LBD, consistent with the X-ray crystal structure of the kainate-bound GluK4 LBD (PDB 5IKB)^60^ (Fig. 4B).

**Figure 4.**
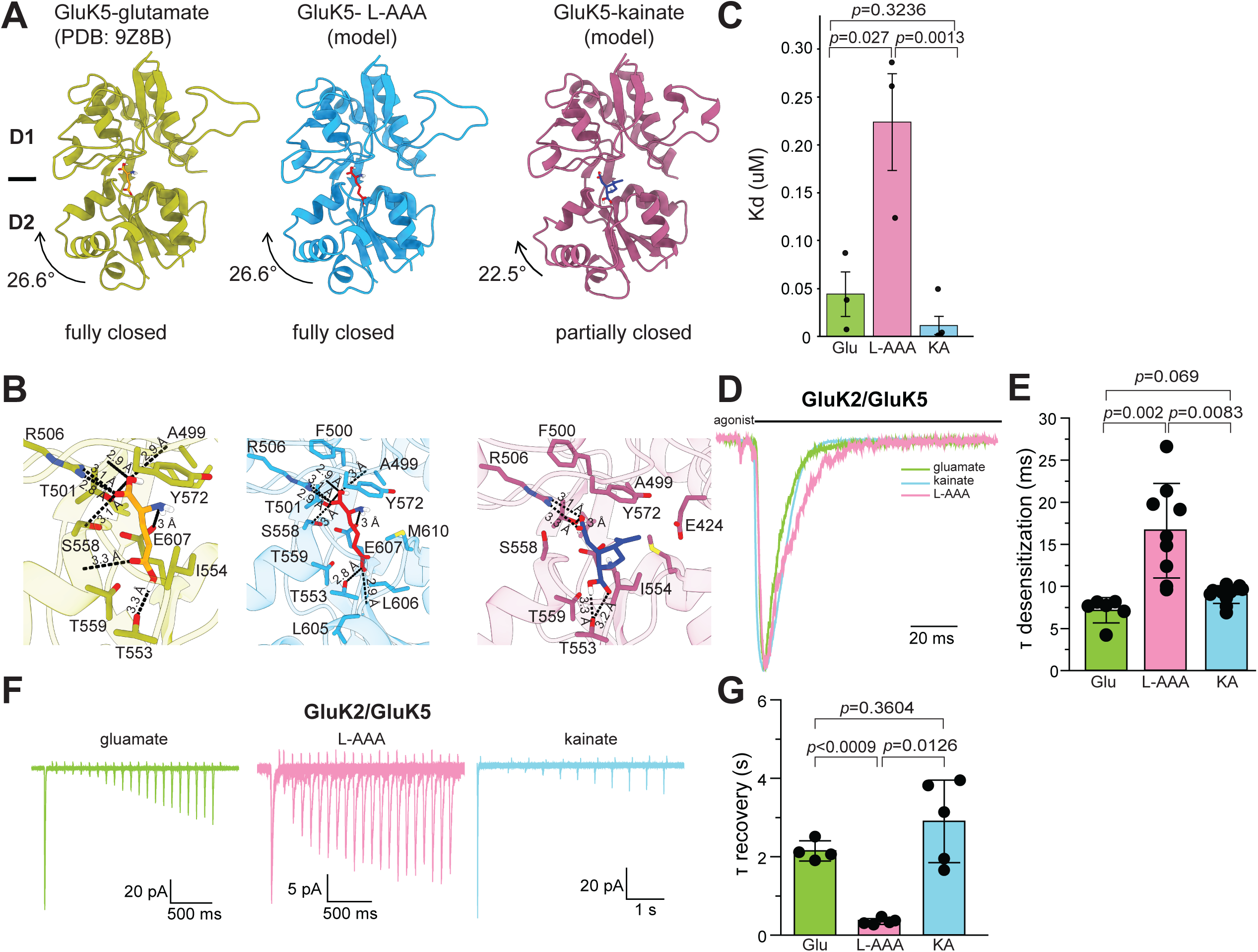
Structural modeling of GluK5 LBD and gating kinetics of GluK2/GluK5 receptors. (**A**) Structural models of the GluK5 LBD in complex with glutamate (PDB: 9Z8B), and computational homology models bound to L-AAA and kainate. D1 and D2 lobes are labeled, with the domain closure angle indicated for each complex relative to the apo state. (**B**) Detailed views of the orthosteric binding pocket for GluK5 bound to ligands. Key residues coordinating the ligands are shown as sticks with dashed lines indicating hydrogen bonds and electrostatic interactions. (**C**) Equilibrium dissociation constants (Kd) for the high-agonist affinity subunit GluK4 measured by MST across glutamate (Glu, green), L-AAA (pink), and kainate (KA, blue). Individual data points represent independent measurements, with bars indicating mean ± SEM. Statistical significance was calculated using student t-test. (**D**) Representative normalized current traces recorded from heteromeric GluK2/GluK5 receptors in response to 1 s application of glutamate, L-AAA, or kainate. The duration of agonist application is indicated by the black bar. (**E**) Summary of desensitization time constants ( *τ*_desensitization_ , ms) for heteromeric GluK2/GluK5 receptors activated by glutamate, L-AAA, and kainate. Individual recordings are shown as black circles, with bars representing mean ± SEM. (**F**) Representative current traces recorded during paired-pulse protocols evaluating recovery from desensitization in GluK2/GluK5 heteromers. (**G**) Summary bar graph comparing recovery time constants (*τ*_recovery_ , s) across three ligands for GluK2/GluK5 heteromers. Individual data points are shown with bars representing mean ± SEM.

To determine Kd, we sought to purify isolated LBDs from high-affinity KAR subunits (GluK4–GluK5). Because rat GluK5 LBD failed to express in Escherichia coli (E.coli), was unstable during expression in insect and mammalian cells and degraded substantially during the expression step and suffering rapid degradation, we used GluK4 LBD instead. To facilitate purification, track expression levels, and enable detection for microscale thermophoresis (MST), constructs were engineered with an N-terminal hexahistidine tag and thrombin cleavage site, alongside a C-terminal fusion containing an HRV 3C protease site, EGFP, and a 1D4 tag. The EGFP fused GluK4 LBD was overexpressed in Sf9 cells and purified via 1D4 affinity chromatography (Fig. S2A) and used for the MST binging analysis. As expected based on the previous reports^60,61^, the Kd of these agonists on the high-agonist-affinity subunit GluK4 are significantly smaller than those on GluK2, with the Kd of kainate (Kd: 16.3 ± 9.5 nM, n=5), glutamate (Kd: 44.0 ± 23.2 nM, n=3), and L-AAA (Kd: 223.5 ± 50.4 μM, n=3) (Fig. 4C). Consistently, L-AAA exhibited a significantly higher Kd value than glutamate and kainate.

### Agonist Desensitization and Recovery Kinetics comparing homo- and heteromeric KARs

To assess how incorporating the high-agonist affinity subunits into the tetrameric KARs influences the desensitization and recovery kinetics triggered by these partial agonists, we co-expressed rat GluK2 and rat GluK5 with 1:3 (w/w) ratio of plasmids in HEK293T cells as previously reported recordings^37,62,63^, and conducted whole-cell patch clamp recordings. Agonist-dependent differences were also observed in the heteromeric GluK2/GluK5 receptors. Desensitization decay kinetics were all fitted with a single exponential function, consistent with observations for heteromeric GluK2. Glutamate and kainate yielded mean τ_desensitization_ values of 7.1 ± 0.6 ms (n=6) 9.0 ± 0.3 ms (n=10), respectively, whereas L-AAA produced significantly slower desensitization in heteromeric receptors (16.6 ± 1.9 ms, n=9) (Fig. 4D, E). When comparing desensitization kinetics between homomeric GluK2 and heteromeric GluK2/GluK5, we observed no statistically significant differences for glutamate or L-AAA activated receptors (p > 0.05). Kainate-activated GluK2/GluK5 heteromers desensitized significantly faster than kainate-activated homomeric GluK2 (p = 0.038), though the absolute difference was minimal. Finally, we evaluated the recovery from desensitization for GluK2/GluK5 heteromers. The recovery time constant (τ_recovery_) for GluK2/GluK5 activated by glutamate, L-AAA, and kainate were 2.1 ± 0.3 s (n=4), 0.4 ± 0.1 s (n=5), and 2.8 ± 1.1 s (n=5), respectively (Fig. 4 F, G). Compared to the τ_recovery_ of homomeric GluK2, recovery of heteromeric KARs was nearly two-fold slower for glutamate (p = 0.00038) and L-AAA (p = 0.0143) but showed no significant difference for kainate (p= 0.667). Considering the structures, we attribute this delayed recovery in heteromeric KARs to the GluK2/GluK5 desensitization ring being stabilized by an expanded network of polar interactions relative to homomeric GluK2^57^, which increases structural stability and impairs resensitization kinetics.

## Discussion

We have previously reported (1) that desensitized homomeric GluK2 KAR adopt a unique conformation that governs the desensitization mechanism markedly distinct from those of other iGluRs, (2) how the bulky partial agonist domoate stabilizes intermediate desensitized conformations of homomeric KARs, and (3) how full agonists and competitive antagonists induce conformational dynamics in heteromeric GluK2/GluK5 KARs that are distinct from those in homomeric GluK2 KARs. Building on these studies, we sought to understand how a variety of partial agonists, spanning a spectrum from near-full agonists to larger, sterically bulky compounds comparable in size to competitive antagonists, dictate desensitization and recovery of KARs, whether through shared mechanisms across homo- and heteromers or via heteromer-specific kinetics.

Leveraging recent advances, we first compared the structures of the GluK2 LBD complexed with five ligands, including the full agonist glutamate, the partial agonists L-AAA, kainate, and domoate, and the competitive antagonist UBP310, derived from experimental structures or computational modelling. The small partial agonist L-AAA is predicted to induce LBD bi-lobe closure to a degree similar to glutamate. As we reported earlier, although domoate is closely comparable in molecular weight and surface area to UBP310, it promotes partial bi-lobed closure similar to that of kainate^46^, whereas UBP310 stabilizes a significantly more open LBD conformation (note that GluK2 was mutated to mimic GluK1 to increase UBP310 affinity)^64^ relative to the apo state^46^ (Fig. 5). The relative binding affinities of these agonists for GluK2 (domoate > kainate > glutamate >> L-AAA) follow the same rank order as published EC_50_ values^45,62^. We observed that desensitization rates followed the rank order glutamate > L-AAA ≈ kainate > domoate (because UBP310 stabilizes the open LBD conformation, inhibits GluK2, and elicits no macroscopic currents, it was excluded from desensitization analysis) (Fig. 5).

**Figure 5.**
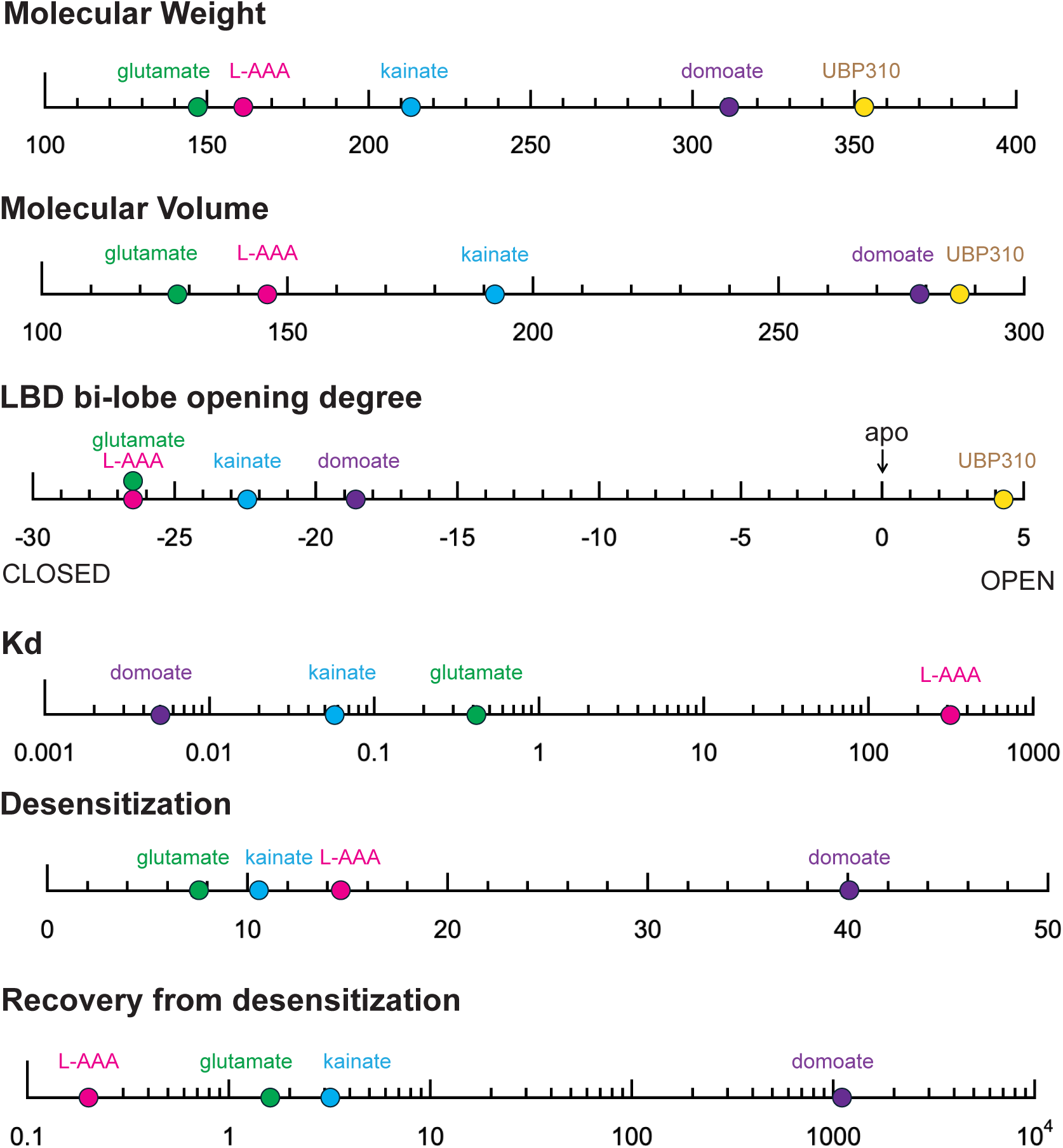
Comparison of biophysical, structural, thermodynamic, and kinetic properties of KAR orthosteric ligands on GluK2 KAR. Linear scale representations comparing select agonists and antagonists (glutamate, L-AAA, kainate, domoate, UBP310, and apo state where applicable) across key chemical and pharmacological parameters.

KAR desensitization is driven by ligand-induced conformational changes, including closure of the LBD bi-lobes, which in turn leads to LBD dimer dissociation and LBD in-plane rotation^1,43,44^. As the full LBD closure induced by glutamate places strong mechanical strain on the LBD dimer interface and triggers rapid separation of the D1–D1 domain interface, which force the receptor quickly into the desensitized state. Across the agonists we tested, the extent of desensitization correlated directly with the degree of LBD bi-lobe closure stabilized by each ligand, except L-AAA. The moderate LBD closure induced by kainate desensitizes receptors rapidly, though not as fast as glutamate, whereas domoate, which stabilizes the least closed LBD state^46^, induces the slowest desensitization. Our computational modelling indicates that L-AAA closes the LBD bi-lobes to an extent similar to glutamate, though the exact degree of closure may vary slightly given the rotational flexibility of the L-AAA carbon backbone, as evidenced by the multiple L-AAA conformations observed in complexes with other proteins, such as aminotransferase^53^ and piperideine-6-carboxylate dehydrogenase^54^. Nevertheless, the main cause of this slower desensitization kinetics observed with L-AAA presumably originated from its rapid unbinding kinetics and low binding affinity. Transition to the desensitized state requires the agonist to remain bound long enough for the LBD to undergo structural rearrangement, including LBD dimer interface rupture and in-plane rotation of the A/C subunits, as described above. Because L-AAA dissociates rapidly, a subset of binding events terminates before the LBD layer transitions into the full desensitized conformation^55^, effectively slowing the macroscopic rate of desensitization entry. However, under continuous application at high concentrations, rapid re-binding maintains high occupancy, allowing the channel population to eventually accumulate almost completely in the thermodynamically favored desensitized state. Indeed, we previously captured a structure of GluK2 in complex with domoate that was stabilized in a transition state, in which one LBD dimer lacked agonist within its binding pockets while the other dimer pair was completely dissociated in the presence of domoate, preventing the receptor from transitioning into a fully desensitized conformation^46^. Although technical limitations prevented us from solving a structure of full-length KAR bound to L-AAA, we propose that L-AAA similarly promotes a transition state by rapidly dissociating from the LBD, thereby slowing or preventing full desensitization.

In addition, our data show that recovery rates of GluK2 activated by these four agonists inversely correlate with Kd. They also correlate with LBD domain closure, except in the case of L-AAA. Recovery from L-AAA induced desensitization is exceptionally fast primarily due to its very high Kd. Although L-AAA is expected to stabilize the LBD closure comparable to glutamate, its rapid off-rate allows the ligand to dissociate rapidly from the desensitized receptors, facilitating a rapid transition back to the resting state. This fast-unbinding mechanism demonstrates how overall desensitization and recovery kinetics are defined by an interplay between the degree of LBD closure and ligand binding/unbinding kinetics. Unlike homomeric GluK1 KARs, which exhibit biphasic recovery, especially those expressing in the hippocampus^3,65–67^, we observed that homomeric GluK2 KARs activated by distinct partial agonists, as well as in our earlier observation with domoate^46^, showed only rapid, single-exponential recovery kinetics, and underwent complete desensitization, except in the presence of domoate^46^.

Next, we investigated whether partial agonists regulate homomeric and heteromeric KARs through similar or distinct mechanisms. Structural modeling revealed that the degree of LBD bi-lobe closure in partial-agonist–bound GluK5 closely resembles that of GluK2, even though the unliganded apo GluK5 LBD is stabilized in a conformation 4° more open than the apo GluK2 LBD^57^. Furthermore, the MST assays demonstrated that the rank order of dissociation kinetics between low-affinity (GluK1–3) and high-affinity (GluK4–5) KAR subunits was conserved across all three agonists, and high-affinity subunits (GluK4–GluK5) displaying approximately an order of magnitude higher sensitivity, consistent with previous reports for glutamate and kainate^68,69^,

While earlier studies showed that desensitization of recombinant GluK2/GluK5 receptors activated by glutamate is faster than that of homomeric GluK2^38,70^, we observed that the desensitization kinetics of heteromeric GluK2/GluK5 receptors activated by saturating or near-saturating concentrations of full and partial agonists were similar to those of homomeric GluK2 KARs. This is consistent with previous observations that the desensitization kinetics of heteromeric KARs are largely agonist-concentration dependent with homomeric and heteromeric KARs behaving similarly at high agonist concentrations^37^. Previously, we utilized a heteromeric trafficking strategy derived from the G-protein-coupled GABA_B_ receptor, which allows only heteromeric receptors to reach the plasma membrane and using this approach, we observed that heteromeric GluK2/GluK5 receptors displayed no significant difference in desensitization kinetics compared to GluK2 in the presence of 10 mM glutamate as well consistent with data from the present study^57^. In the presence of high concentration of agonist, this observation aligns with the proposed model wherein high-affinity subunits drive receptor activation, whereas low-affinity subunits (GluK1–GluK3) control desensitization, a conserved behavior we observed across both full and partial agonists. As Khanra et al. and our group have previously observed^34,35,57^, the GluK2 subunits at the BD positions in homotetrameric GluK2 predominantly undergo conformational changes to drive assembly of the desensitization ring, whereas the remaining two subunits at the AC positions undergo minimal movement upon desensitization. Consequently, replacing the AC subunits with GluK5 in heterotetramers does not appear to drastically alter the fundamental energy barriers governing recovery from desensitization. Consistent with these structural features, our data show that LBD dimer dissociation induced by various partial agonists to initiate desensitization does not differ between homomeric and heteromeric subtypes.

In summary, our data show that the intrinsic pharmacological properties of agonists predominate over subunit composition in defining KAR desensitization.

## Materials and Methods

### Plasmid construction

For electrophysiological experiments, rat wild-type GluK2 and GluK5 were subcloned into pCAG/IRES/EGFP (Addgene #119739) and pCAG/IRES/mCherry (Addgene #45766) vectors, respectively. For KAR LBD protein expression, LBD constructs were generated based on a previously published SUMO-fused GluN2A LBD construct^71^. The S1 and S2 domains of rat GluK2 were connected via a GT linker, fused to an N-terminal hexa-His tag and SUMO tag, and subcloned into the pET-22b vector (synthesized by and obtained from GenScript) for bacterial expression. Similarly, the S1 and S2 domains of rat GluK4 were connected via a GT linker and fused to an N-terminal octa-His tag, and a C-terminal HRV 3C protease cleavage site, EGFP, and a 1D4 tag. This construct was subcloned into a pFp10 vector containing HR1 and HSP70 promoters for insect expression^72,73^.

### Cell culture

For electrophysiology experiments, human embryonic kidney 293T (HEK293T) cells (ATCC, Cat. No. CRL-3216) were cultured in Dulbecco’s modified Eagle medium (DMEM, Corning) supplemented with 10% FBS at 37°C in a 95% O₂–5% CO₂ atmosphere. HEK293T cells were transfected using the TransIT2020 transfection reagent (Mirus) according to the manufacturer’s instructions. At 18-24 hours post-transfection, cells were dissociated using Accutase (Innovative Cell Technologies, Inc.), resuspended, and plated onto 35 mm poly-D-lysine-coated dishes (Neuvitro). Electrophysiological recordings were performed 2–4 hours later^74^. For biophysical analysis, the GluK2 LBD were transformed in Origami 2 (Novagen) which grown in Lysogeny Broth (LB) medium. Cultures were adjusted to an initial optical density at 600 nm (OD_600_) of 0.1 and incubated at 37 °C with shaking at 200 rpm for approximately 8 hours until reaching an OD_600_ of 1.5-2.0. The culture flasks were transferred to 4 °C for 30 minutes to cool down prior to induction. Protein expression was induced by the addition of isopropyl *β* -D-1-thiogalactopyranoside (IPTG) to a final concentration of 0.8 mM, and cultures were incubated for 20hrs at 15 °C . Cells were harvested and stored for subsequent protein purification. Baculovirus containing the GluK4 LBD construct was prepared using standard protocols. Protein expression was carried out in Sf9 cells incubated at 27 °C for 48 hours. Expression cultures were harvested by centrifugation, and cell pellets were stored at −80 °C until further use.

### Electrophysiological recordings

All whole-cell recordings were performed using HEKA EPC10 amplifiers (HEKA Elektronik, Lambrecht, Germany) with thin-wall borosilicate glass pipettes (2–5 MΩ) coated with dental wax to reduce electrical noise. Currents were recorded at a holding potential of −70 mV, with a sampling frequency of 10 kHz, and filtered at 2.6 kHz. The external solution contained (in mM): 145 NaCl, 2.5 KCl, 1.8 CaCl₂, 1 MgCl₂, 5 glucose, and 5 HEPES. The internal pipette solution contained (in mM): 105 NaCl, 20 NaF, 5 Na₄BAPTA, 0.5 CaCl₂, 10 Na₂ATP, and 5 HEPES. The pH and osmotic pressure of the external and internal solutions were adjusted to 7.4 and 300–290 mOsm/kg, respectively. L-glutamate was applied using theta glass tubing mounted on a piezoelectric stack (MXPZT-300 series, Siskiyou) driven by a HEKA EPC10 amplifier. Typical 10– 90% rise times were 250–300 µs, measured from junction potentials at the open tip of the patch pipette after recordings. Data acquisition was performed using PULSE software (HEKA Elektronik, Lambrecht, Germany).

### Electrophysiological data analysis

The macroscopic rate of desensitization (τ_desensitization_) was measured by fitting an exponential function to the decay of current from ∼80% of its peak amplitude (Ipeak) to baseline in recordings where glutamate was applied for 1 second. Desensitization kinetics were fitted using a single-exponential, one-term Levenberg-Marquardt fitting approach. To evaluate recovery from desensitization, two protocols were employed: (1) a two-pulse application of 10 mM glutamate for 100 ms per pulse, with interpulse intervals increasing in 100 ms steps from 50 ms to 2 s; and (2) a second protocol with interpulse intervals increasing in 2 s steps, ranging from 50 ms to 20 s. In all cases, the first peak was considered the control (100% activation), while the second peak represented recovery at a given time point. Data from both protocols were combined, and a recovery curve was generated, plotting the percentage of recovery against time. The experimental data were fitted to one-term exponential model, from which the recovery time constant (τ) was derived. Statistical analysis. Because group variances and sample sizes were unequal, we used Welch’s ANOVA with Games–Howell post-hoc comparisons (two-sided, α = 0.05). Exact P values are reported. Analyses were performed in OriginPro 2025.

### Ligand structure selection and physicochemical descriptor analysis

The molecular and physicochemical properties of L-glutamate, L-AAA, kainic acid, domoic acid, and UBP310 were compared using standardized stereochemically defined structures obtained from PubChem. The PubChem compound identifiers used were CID 33032 for L-glutamate, CID 92136 for L-α-aminoadipic acid, CID 10255 for kainic acid, CID 5282253 for domoic acid, and CID 6420160 for UBP310. Structures were inspected to verify compound identity, stereochemistry, and the absence of counterions, protecting groups, or covalent derivatives. For L-α-aminoadipic acid, the stereochemically defined (2S) compound, CID 92136, was used. Records corresponding to compounds with undefined stereochemistry, the (2R) enantiomer, salts, protected derivatives, or structurally related analogues were excluded.

Molecular descriptors were calculated using the JavaScript implementation of RDKit (version 2025.3.4-1.0.0). Molecular weight and exact mass were calculated from the neutral molecular structure. Topological polar surface area (TPSA), number of rotatable bonds, hydrogen-bond donors and acceptors, and heavy-atom count were calculated from explicitly defined SMILES representations of the predominant physiological microstates at pH 7.4. Physiological microstates were assigned based on the expected ionization of carboxylic acid and aliphatic amino groups at pH 7.4. Carboxylic acid groups were represented as deprotonated carboxylates, whereas aliphatic amino groups were represented as protonated. The resulting formal net charges were −1 for L-glutamate, −1 for L-AAA, −1 for kainate, −2 for domoate, and −1 for UBP310. These assignments represent chemically inferred predominant microstates and were not derived from experimental or computational pKa determinations. Three-dimensional shape descriptors were calculated from the default PubChem3D conformer of each stereochemically defined compound. Molecular volume was estimated as the union of atomic van der Waals spheres using a three-dimensional grid with a spacing of 0.20 Å. Bondi-like van der Waals radii of 1.20, 1.70, 1.55, 1.52, and 1.80 Å were used for H, C, N, O, and S atoms, respectively.

Solvent-accessible surface area (SASA) was calculated using a custom implementation of the Shrake–Rupley algorithm, with a solvent-probe radius of 1.40 Å and 960 approximately uniformly distributed surface points per atom. Molecular maximum dimension was calculated as the largest interatomic distance in each conformer after adding the van der Waals radii of the corresponding atom pair. Calculations were performed using Python 3.12.13 and NumPy 2.3.5. The PubChem3D coordinates corresponded to neutral-form conformers and were not independently minimized after assigning the physiological protonation states. Therefore, molecular volume, SASA, and maximum dimension were interpreted as standardized single-conformer shape descriptors rather than as ensemble-averaged properties of the predominant ionic species in solution. All ligands were analyzed using identical atomic radii and numerical parameters.

### Recombinant Protein Purification and Binding Analysis

#### *Purification of GluK2 LBD:* Cell pellets obtained from a 6 L bacterial culture were resuspended in

240 mL of lysis buffer comprising 20 mM HEPES (pH 7.3), 200 mM NaCl, and 1 mM phenylmethylsulfonyl fluoride (PMSF). Resuspended cells were lysed using a sonicator. The lysate was clarified by ultracentrifugation at 40,000 rpm for 40 min at 4°C, and the resulting supernatant was collected for purification. The clarified soluble supernatant was loaded onto the Ni-NTA beads (Cat. No. 17-0575-02, GE Healthcare) equilibrated with a buffer containing 20 mM HEPES (pH 7.3), 200 mM NaCl, and 20 mM imidazole. Unbound and non-specifically bound proteins were washed from the column using sequential step washes with 20 and 40 mM imidazole. Bound his tagged GluK2 LBD was eluted with ∼250 mM imidazole. Peak fractions were collected and used for binding analysis.

#### Purification of GluK5 LBD

Cell pellets harvested from approximately 2 L of Spodoptera frugiperda (Sf9) culture were resuspended in lysis buffer containing 20 mM HEPES (pH 7.3), 200 mM NaCl, and 1 mM PMSF. Resuspended cells were lysed by sonication. The lysate was clarified by ultracentrifugation at 40,000 rpm. The resulting supernatant containing GluK4 LBD was collected for affinity purification. Strep-tactin resin (Cat No. 2-5030-025, IBA) was equilibrated and incubated with the clarified soluble supernatant, and the resin–lysate suspension was then packed into a gravity-flow column. The resin was washed extensively with wash buffer 1 (20 mM HEPES pH 7.3, 150 mM NaCl), wash buffer 2 (20 mM HEPES pH 7.3, 150 mM NaCl, 3 mM Mg-ATP), wash buffer 3 (20 mM HEPES pH 7.3, 500 mM NaCl), and wash buffer 4 (20 mM HEPES pH 7.3, 150 mM NaCl). GluK4 LBD was eluted with an elution buffer containing 20 mM HEPES (pH 7.3), 150 mM NaCl, and 50 mM biotin. To remove the EGFP-Strep fusion tag, the eluted protein was subjected to 3C protease cleavage. Recombinant Human Rhinovirus 3C protease was added to the protein sample at a mass ratio of 1:2 (3C protease to GluK4 LBD) and incubated for 1 h at 4°C. Following digestion, the reaction mixture was applied to pre-equilibrated Streptactin resin and the flow-through containing the cleaved, tag-free protein target was collected and used for the binding assay.

#### MST recording

His-tagged LBDs were labeled with fluorescent dye using His-Tag Labeling Kit RED-tris-NTA 2nd Generation (Cat No. MO-L018, Nanotemper Technologies) in accordance with the manufacturer’s instructions. All dye-containing stock solutions, reaction mixtures, and resulting fluorophore-labeled protein conjugates were protected from ambient light exposure throughout all handling, incubation, and storage procedures. Ligand-binding affinities (Kd) of fluorescently labeled His-tagged LBDs were measured using MicroScale Thermophoresis (MST) on a Monolith NT.115 instrument (NanoTemper Technologies). A 16-point 1:1 serial dilution series of the ligand was prepared in phosphate-buffered saline containing 0.05% Tween-20. Standard Premium Capillaries (Cat No MO-K025, Nanotemper Technologies) were filled with reaction mixture, and Measurements were conducted on a Monolith NT instrument using the RED dye channel setup under Excitation LED power 80–85%, and MST power Medium. Data Processing: Experiments were performed in triplicate. Normalized fluorescence values or raw MST traces were plotted against ligand concentration and fitted to a 1:1 binding model using the Monolith Analysis software to calculate the Kd.

### Biophysical Analysis

The purified His-tagged proteins were dialyzed against HBS buffer containing 20 mM HEPES (pH 7.3) and 150 mM NaCl, then labeled using a His-tag labeling kit (NanoTemper Technologies, cat. no. MO-L018) according to the manufacturer’s protocol. Protein concentrations were determined using a NanoDrop spectrophotometer. A 16-point serial dilution series was prepared, and all measurements were performed in triplicate.

## Key source table

| **Reagent type (species) or resource** | **Designation** | **Source or reference** | **Identifiers** | **Additional information** |
|---|---|---|---|---|
| Gene (*Rattus norvegicus*) | GRIK2_RAT | Genscript (synthesized) | P42260 |  |
| Chemical compound, drug | L-glutamate | Sigma Aldrich | Cat. No. 49621 |  |
| Chemical compound, drug | Kainate | Sigma-Aldrich | Cat.No:K0250 |  |
| Chemical compound, drug | L-alph-aminoadipatic acid | Sigma-Aldrich | Cat. No: A7275 |  |
| Software, algorithm | Pulse | HEKA | https://scicrunch.org/resolver/SCR_018399 | https://www.heka.com/ |
| Software, algorithm | OriginPro 2020 | OriginLab | RRID:SCR_014212 | https://www.originlab.com/2020 |
| Software, algorithm | SWISS-MODEL | Expasy |  | https://swissmodel.expasy.org/interactive |
| Software, algorithm | Autodock vina | doi:10.1038/nmeth.4169. | RRID:SCR_016501 | https://cryosparc.com/ |
| Software, algorithm | UCSF Chimera | doi:10.1002/jcc.20084 | RRID:SCR_004097 | http://plato.cgl.ucsf.edu/chimera/ |

## Acknowledgment

We are grateful to Drs. Corey Smith and Shyue-An Chan at CWRU for their technical assistance with patch-clamp electrophysiology. This work was funded by the National Institute of Health (1R35GM147266-01 to N.T.).

## Author contributions

S.G., and N.T. conceived the project and designed the experimental procedures. C.Z. analyzed structures and models. S.G. conducted electrophysiological recordings. A.H. conducted binding assays. C.Z., S.G., and N.T. wrote the manuscript. All authors reviewed the final draft.

---

**Figure S1. Structural analysis of L-AAA binding modes, molecular dimensions, and alignment within the GluK2 LBD.**

**(A)** Structural overlay of L-α-aminoadipic acid (L-AAA) binding conformations from PDB entries 6HNV (purple) and 6RTU (grey), highlighting flexibility in the side-chain backbone alignment. **(B)** Correlation plot of solvent-accessible surface area versus molecular weight for orthosteric ligands: glutamate (green), L-AAA (magenta), kainate (blue), domoate (purple), and UBP310 (yellow). **(C)** Structural superposition of the glutamate and L-AAA bound GluK2 LBD highlighting the shared the LBD bi-lobe arrangement. **(D)** Close-up structural overlay comparing bound glutamate (green, PDB: 1S50) with the computational homology model of L-AAA bound to GluK2 LBD (magenta). **(E)** Comparison between the crystal structure conformation of L-AAA from PDB 6HNV (purple) and the modeled binding conformation of L-AAA in GluK2 LBD (magenta).

**Figure S2. Expression of high-affinity subunit, structural alignment between low and high-affinity subunits, and primary sequence conservation within the ligand-binding pocket.**

**(A)** Biochemical characterization of purified high-affinity KAR subunit GluK4 LBD construct. SDS-PAGE with Coomassie blue staining (left) and EGFP-fluorescent imaging (right) demonstrating high purity and correct molecular weight. **(B)** Structural superposition of homology models of L-AAA bound GluK5 LBD (dark blue) and GluK2 LBD (light purple). Both domains adopt a conserved clamshell in the presence of L-AAA. **(C)** Multiple sequence alignment of key D1 (top) and D2 (bottom) lobe residues across KAR subunits (GluK1–GluK5), showing the conserved residues in the ligand binding pockets.

