## supplemental figures for "Partial agonists and heteromeric Kainate receptor desensitization"

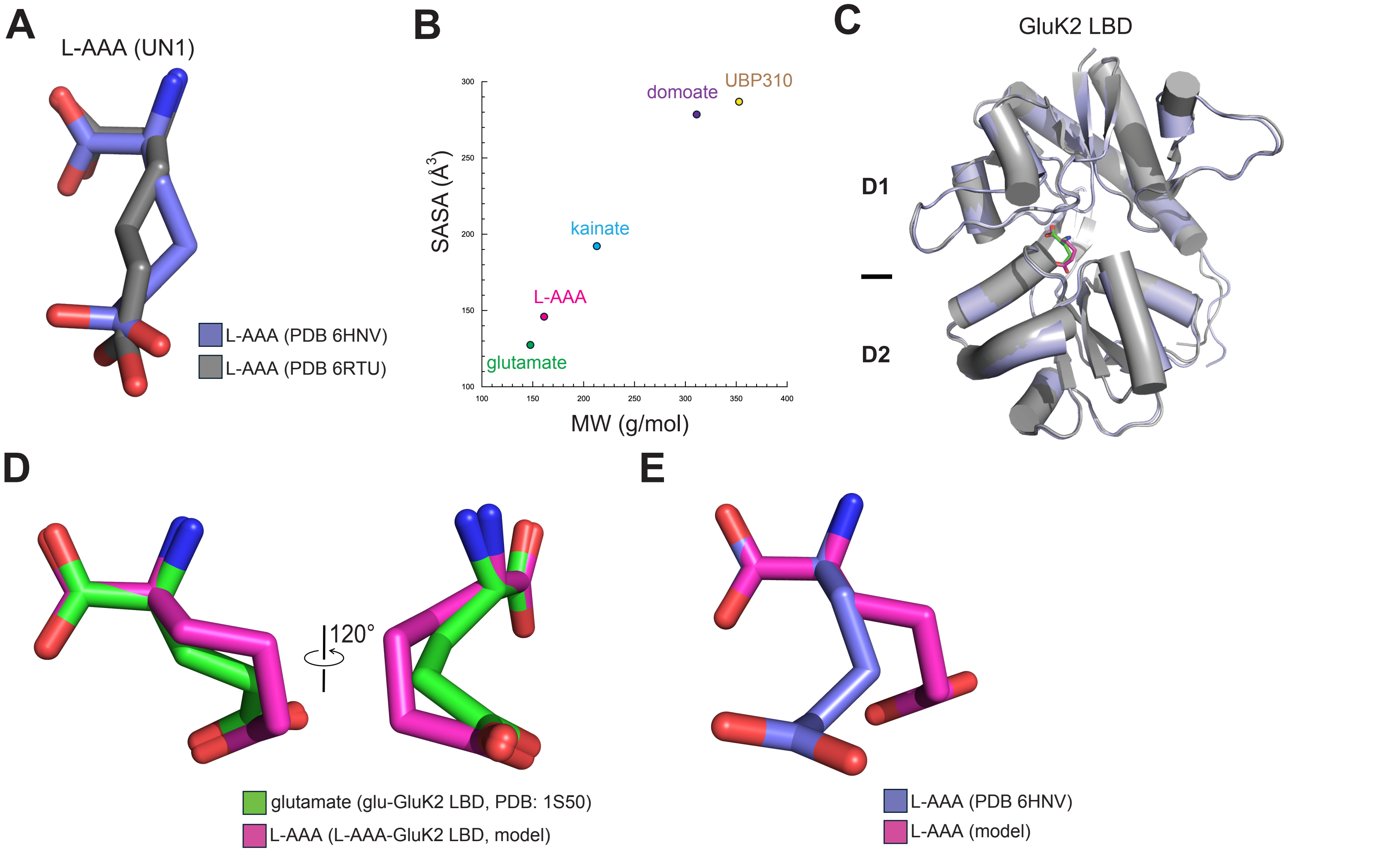
A
L-AAA (UN1)
L-AAA (PDB 6HNV)
L-AAA (PDB 6RTU)
B
SASA (Å³)
MW (g/mol)
300
250
200
150
100
100
150
200
250
300
350
400
domoate
UBP310
kainate
L-AAA
glutamate
C
GluK2 LBD
D1
D2
D
120°
glutamate (glu-GluK2 LBD, PDB: 1S50)
L-AAA (L-AAA-GluK2 LBD, model)
E
L-AAA (PDB 6HNV)
L-AAA (model)

### Figure S1

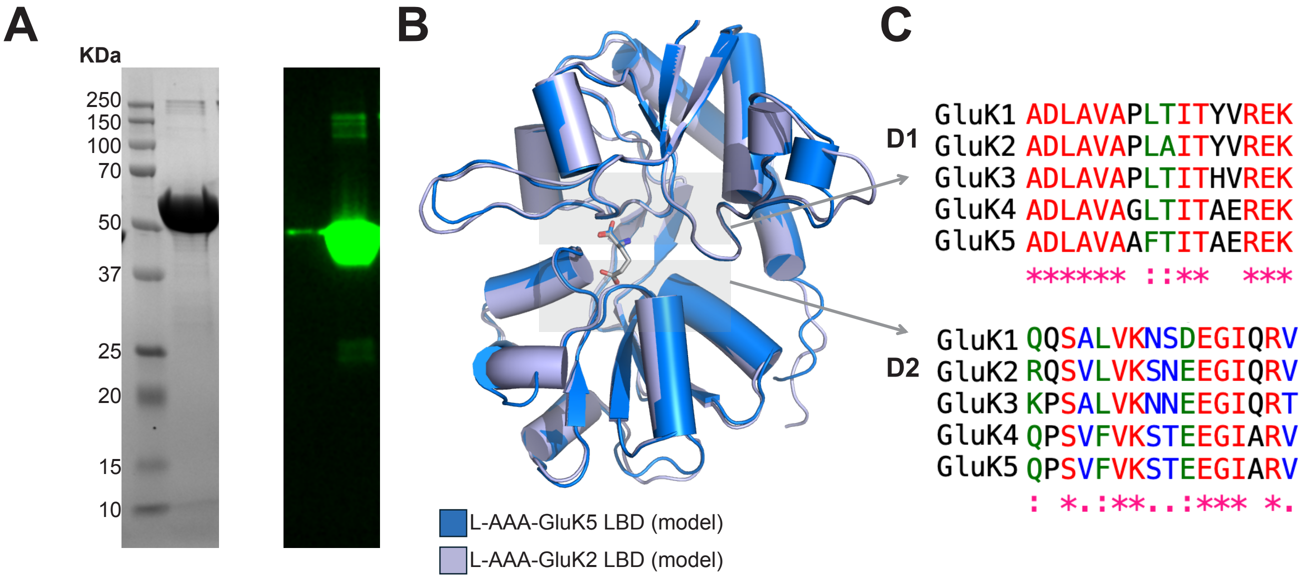
A
KDa
250
150
100
70
50
37
25
20
15
10
B
L-AAA-GluK5 LBD (model)
L-AAA-GluK2 LBD (model)
C
D1
GluK1 ADLAVAPLTITYVREK
GluK2 ADLAVAPLAITYVREK
GluK3 ADLAVAPLTITHVREK
GluK4 ADLAVAGLTITAEREK
GluK5 ADLAVAAFTITAEREK
****** ::**  ***
D2
GluK1 QQSALVKNSDEGIQRV
GluK2 RQSVLVKSNEEGIQRV
GluK3 KPSALVKNNEEGIQRT
GluK4 QPSVFVKSTEEGIARV
GluK5 QPSVFVKSTEEGIARV
: *.:**..:*** *.

### Figure S2
